# Platelet pannexin-1 channels modulate inflammation during abdominal aortic aneurysm formation

**DOI:** 10.1101/2022.11.23.517652

**Authors:** Lisa Maria Metz, Tobias Feige, Larissa de Biasi, Agnes Ehrenberg, Joscha Mulorz, Laura Mara Toska, Friedrich Reusswig, Christine Quast, Norbert Gerdes, Malte Kelm, Hubert Schelzig, Margitta Elvers

## Abstract

Abdominal aortic aneurysm (AAA) is a common disease and highly lethal if untreated. The progressive dilatation of the abdominal aorta is accompanied by degradation and remodeling of the vessel wall due to chronic inflammation. Pannexins represent anion-selective channels and play a crucial role in non-vesicular ATP release to amplify paracrine signaling in cells. Thus, pannexins are involved in many (patho-) physiological processes. Recently, Panx1 channels were identified to be significantly involved in AAA formation through endothelial derived Panx1 regulated inflammation and aortic remodeling. In platelets, Panx1 becomes activated following activation of glycoprotein (GP)VI. Since platelets play a role in cardiovascular diseases including AAA, we analyzed the contribution of platelet Panx1 in the progression of AAA. We detected enhanced Panx1 plasma levels in AAA patients. In experimental AAA using the pancreatic porcine elastase (PPE) mouse model, a major contribution of platelet Panx1 channels in platelet activation, pro-coagulant activity of platelets and platelet-mediated inflammation has been detected. In detail, platelets are important for the migration of neutrophils into the aortic wall induced by direct cell interaction and by activation of endothelial cells. Decreased platelet activation and inflammation did not affect ECM remodeling or wall thickness in platelet-specific Panx1 knock-out mice following PPE surgery. Thus, aortic diameter expansion at different time points after elastase infusion of the aortic wall was unaltered in platelet-specific Panx1 deficient mice suggesting that the modulation of inflammation alone does not affect AAA formation and progression. In conclusion, our data strongly supports the role of platelets in inflammatory responses in AAA via Panx1 channels and adds important knowledge about the significance of platelets in AAA pathology important for the establishment of an anti-platelet therapy for AAA patients.

## 1 Introduction

Abdominal aortic aneurysm (AAA) is defined as a permanent dilatation of the abdominal aorta to 1.5 fold to its normal diameter and affects all three layers of the vascular tunica (1, 2). In most cases, AAAs are asymptomatic and frequently incidental findings or detected during systemic screening of populations at risk such as men older than 65 years or people with positive family history (3). However, several risk factors are associated with AAA such as tobacco smoking, advanced age, hypertension, chronic obstructive pulmonary disease, hyperlipidemia, genetic susceptibility and male sex (4). During AAA progression, the aortic wall becomes thinner due to proteolytic degradation of the ECM proteins elastin and collagen over time. Furthermore, AAA pathology is characterized by immune cell infiltration, vascular smooth muscle cell (VSMCs) apoptosis, increased oxidative stress in the aortic wall and the formation of an intraluminal thrombus (ILT) (5, 6). The importance of inflammatory processes in AAA suggest that leukocytes, endothelial cells and platelets might be involved in the initiation and progression of AAA (7).

Pannexins represent a group of transmembrane proteins and function as an ion channel for small molecules up to 1kDa in size (8). Panx1 is a single membrane channel consisting of a hexameric structure of single membrane channels at the plasma membrane. Opening of Panx1 channels is regulated via different mechanisms such as phosphorylation of Src family kinases (SFKs) and elevation of [Ca^2+^]_int_ (8, 9). Panx1 are anion-selective channels with a high conductance for ATP and display an important role in non-vesicular ATP release and intercellular communication by amplification of paracrine signaling in cells (8). Thus, Panx1 channels play an important role in various physiological functions as normal development of skin and bones, synaptic plasticity and the regulation of blood pressure (10, 11). Recent studies demonstrate that Panx1 channels contribute to AAA formation. Increased Panx1 expression levels in aortic tissue have been observed in male AAA patients. In a mouse model of experimental AAA, endothelial derived Panx1 regulated signaling events to mediate inflammation and aortic remodeling (12).

In platelets, Panx1 channels are exposed at the plasma membrane to release ATP upon platelet stimulation with thrombin, collagen or the TxA_2_ analogue U46619 (13, 14). Platelet activation leads to increased Panx1 phosphorylation at Tyr^198^ and Tyr^308^ via the Src-GPVI signaling axes to support arterial thrombus formation (13). Clinical and experimental data suggest that platelets play a pivotal role in AAA pathology because impaired endothelial function causes platelet activation and promotes a persistent renewal of cellular activity at the luminal interface of the ILT via entrapment and recruitment of activated platelets to establish a chronic inflammatory response. However, the specific role of platelets in AAA disease and the impact of platelet Panx1 in AAA progression is not well defined to date.

## 2 Materials and methods

### 2.1 Chemicals and antibodies

For platelet isolation apyrase (Grade II from potato, #A7646, Sigma-Aldrich) and prostacyclin (#P5515, Calbiochem,) were used. Platelets were activated with collagen-related peptide (CRP; Richard Farndale, University of Cambridge, Cambridge, UK), adenosine diphosphate (ADP; #A2754, Sigma-Aldrich), the thromboxane A2 analogue U46619 (U46; #1932, Tocris), PAR4 peptide (PAR4; St. Louis, Missouri, MO, USA), thrombin (Thr; #10602400001, Roche Diagnostics, Germany), lipopolysaccharide (LPS, #L4524, Sigma-Aldrich) or tumor necrosis factor alpha (TNFα, #300-01A, Preprotech). For immunoblotting antibodies targeting Pannexin-1 (Panx1, #91137S, Cell Signaling, 1:1000) and phosphorylated Panx1 (pPanx1 Tyr^198^, #ABN1681, Merck, 1:1000), glyceraldehyde 3-phosphate dehydrogenase (GAPDH, #2118S, Cell Signaling, 1:1000) and horseradish peroxidase (HRP)-linked anti-rabbit secondary antibodies (#7074S, Cell Signaling, 1:2500) were used. For flow cytometric analysis fluorophore conjugated antibodies labelling P-selectin (CD62P, Wug.E9-FITC, #D200, Emfret Analytics, 1:10), active integrin αIIbβ3 (JON/A-PE, #D200; Emfret Analytics, 1:10), GPIbα (CD42b, Xia.G5-PE, #M040-2 Emfret Analytics, 1:10), GPVI (JAQ1-FITC, #M011-1, Emfret Analytics, 1:10), integrin α5 (CD49e, Tap.A12-FITC, # M080-1, Emfret Analytics, 1:10), integrin β3 (CD61, Luc.H11-FITC, #M031-1, Emfret Analytics, 1:10) Ly6G (1A8-APC, #127614, BD Biosciences, 1:30) and phosphatidylserine (PS) exposure (CyTM 5 Annexin V, #559934, BD Bioseciences, 1:10) were used. Immunofluorescence staining was performed using protein blocking solution (#X0909, Dako) antibodies against GPIbα (CD42b, #M042-0, Emfret Analytics, 1:50) and Ly6G (#551459; BD Pharming, 1:100), biotinylated secondary antibodies (#BA-9400, Vector, 1:200), streptavidine FluorTM 660 conjugates (#50-4317-80, Thermo Fisher Scientific, 1:20) and DAPI (#10236276001, Roche, 1:3000) was used. MHEC5-T cells were cultivated in DMEM high glucose cell culture medium (#41965062, Gibco).

### 2.2 Human blood samples and ethic votes

Fresh citrate-anticoagulated blood (BD-Vacutainer®; Becton, Dickinson and Company; #367714) was collected from abdominal aortic aneurysm (AAA) patients before open surgery or endovascular aortic repair (EVAR). Blood samples of healthy volunteers with an age ≥ 60 years served as age-matched controls (AMCs). All AAA patients and healthy volunteers provided their written informed consent to participate in this study according to the Ethics Committee of the University Clinic of Duesseldorf, Germany (2018-140-kFogU, study number: 2018064710; biobank study number: 2018-222_1 and MELENA study: 2018-248-FmB, study number: 2018114854). This study was conducted according to the Declaration of Helsinki and the International Council for Harmonization Guidelines on Good Clinical Practice. The procedure of this study was also approved by the Ethics Committee of the University Clinic of Duesseldorf, Germany.

### 2.3 Human platelet isolation and plasma preparation

Citrate-anticoagulated blood (BD-Vacutainer®; Becton, Dickinson and Company; #367714) was centrifuged at 200x g for 10 min at room temperature (RT). After centrifugation, the platelet-rich plasma (PRP) was collected and added to PBS pH 6.5 (supplemented with 2.5 U/mL apyrase and 1 μM PGI_2_) in a volumetric ratio of 1:1, followed by a centrifugation step at 1000x g for 6 min. The platelet pellet was resuspended in Tyrode’s buffer pH 7.4 (containing 140 mM NaCl_2_, 2.8 mM KCl, 12 mM NaHCO_3_, 0.5 mM Na_2_HPO_4_, 5.5 mM Glucose). The platelet cell number was measured using a hematology analyzer (Sysmex - KX21N, Norderstedt, Germany) and adjusted for the following experiments. For human plasma preparation, fresh citrate-anticoagulated blood was centrifuged for 10 min at 1500x g at 4 °C. The platelet free plasma was collected and stored at −70 °C until use.

### 2.4 Human platelet lysates and immunoblotting

Human platelets were isolated as described in 4.3. Isolated platelets (40 × 10^6^) were stimulated with ADP (10 μM), respectively CRP (0.1 or 1 μg/mL) for 10 min at 37 ºC. Platelet stimulation was terminated by centrifugation at 800x g at 4°C for 5 min. Platelet releasates were collected and analyzed for soluble Panx1 via ELISA. The platelet pellet was lysed by adding 1x human lysis buffer (100 mM Tris-HCl, 725 mM NaCl_2_, 20 mM EDTA, 5 % TritonX-100, complete protease inhibitor). Lysis was performed for 15 min at 4 °C. Afterwards, 6x Laemmli buffer was added to the platelet lysates, followed by a denaturation step at 95°C for 5 min. The platelet lysates were stored at −70 °C until use. For immunoblotting the platelet lysates were separated by a SDS-polyacrylamide gel electrophoresis (12 % SDS-polyacrylamide gel) and electro-transferred onto a nitrocellulose blotting membrane (GE Healthcare Life sciences). After blotting, the membrane was blocked with 5 % non-fat dry milk in TBST (Tris-buffered saline, containing 0.1% Tween20) for 1h. The membrane was incubated with primary antibodies against Panx1 (#91137S, Cell Signaling, 1:1000), respectively phospho Panx1 Tyr^198^ (#ABN1681, Merck, 1:1000) at 4°C overnight and with an appropriate HRP-conjugated secondary antibody (#7074S, Cell Signaling, 1:2500) for 1h at RT. GAPDH (#2118S, Cell Signaling, 1:1000) served as loading control. Bands were visualized by chemiluminescence detection reagents (BioRad, #1705061) and imaged using a FusionFX Chemiluminescence Imager System (Vilber). Quantification of immune-reactive band intensities was performed by using the FUSION FX7 software (Vilber).

### 2.5 Animals

Pathogen-free *Panx1*^*fl/fl*^ mice were provided by Dr. Brant Isakson (University of Virginia, Charlottesville, VA, USA) and cross-bread with PF4-Cre mice, that were purchased from the Jackson Laboratory (C57BL/6-Tg [Pf4-cre] Q3Rsko/J), to generate megakaryocyte/platelet specific deletion of Panx1 as described earlier (13).

The PPE surgery was performed only in male mice aged 10-12 weeks. All other experiments were conducted with male and female mice with an age of 2-4 months. Mice were maintained in an environmentally controlled room at 22 ± 1 °C with a 12 h day-night cycle. Mice were housed in Macrolon cages type III with *ad libitum* access to food (standard chow diet) and water. All animal experiments were conducted according to the Declaration of Helsinki and approved by the Ethics Committee of the State Ministry of Agriculture, Nutrition and Forestry State of North Rhine-Westphalia, Germany (Reference number: AZ 81-02.05.40.21.041; AZ 81-02.4.2018.A409).

### 2.6 The porcine pancreatic elastase (PPE) perfusion mouse model

The porcine pancreatic elastase (PPE) perfusion model represents an experimental in vivo model to investigate AAA formation and progression in mice by the infusion of porcine pancreatic elastase into the abdominal aorta. Therefore, mice were anesthetized with 2-3 % isoflurane. Additionally, all mice received a locally subcutaneous (s.c.) injection of buprenorphine [0.1 mg/kg body weight] 3 min prior surgery. The anesthesia was continuously monitored throughout the whole procedure. After laparotomy, an aortotomy above the iliac bifurcation was conducted following the temporal ligation of the proximal and distal infrarenal aorta. Afterwards a catheter was inserted at the distal part of the previously conducted aortotomy, to perfuse the infrarenal aortic segment for 5 min with sterile isotonic saline (NaCl_2_, 0.9 %) containing type I porcine pancreatic elastase (2.5 U/mL, Sigma-Aldrich) at 120 mmHg. After perfusion, the catheter was removed and the aortotomy was closed. After surgery all mice received subcutaneous injection of buprenorphine (0.1 mg/kg body weight) every 6 h within the daylight phase for the following 3 days. Additionally, all operated mice received buprenorphine (0.3 μg/mL in H2O) in the drinking water for a time period of 3 days after surgery. Animals were euthanized to collect aortic tissue and blood samples at day 3, 7, 14, respectively 28 after PPE surgery. The PPE surgeries were conducted by Dr. med. Joscha Mulorz (Department of Vascular- and Endovascular Surgery, University Hospital Duesseldorf) and Julia Odendahl (Department of Cardiology, Pulmonology and Vascular Medicine, University Hospital Duesseldorf; S1-Project TRR259).

### 2.7 Ultrasound imaging of the abdominal aorta

For monitoring of AAA development and progression after PPE surgery, the dilation of the abdominal aorta was analyzed by measuring the aortic diameter within the aneurysm segment via Ultrasound. Ultrasound measurements were performed prior surgery (baseline) and at day 3, 7, 14, 21 and 28 after PPE. For ultrasound imaging, all mice were anesthetized with 2-3 % isoflurane and positioned on a heating plate at 37 °C. All ultrasound measurements were conducted using a Vevo 2100® High-Resolution In Vivo Micro-Imaging System (VisualSonics). Inner aortic diameter and aortic wall thickness were measured using a standardized imaging algorithm with longitudinal B-Mode imaging during the systolic phase.

### 2.8 Preparation of mouse tissue

Mice were kept under constant anesthesia with isoflurane and were euthanized by terminal heart puncture followed by cervical dislocation. For platelet and plasma preparation blood samples were collected. After opening the thorax, the vascular system was rinsed with cold heparin solution (20 U/mL; 4°C) to avoid clot formation. Therefore, the heart was punctured with a butterfly cannula at the apex cordis of the left ventricle. The right atrium was incised and approximately 20 mL of heparin solution were perfused under constant pressure through the vascular system. Afterwards organs, including the thoracic and abdominal aorta, were removed and fixed for 24 h in 4% paraformaldehyde (PFA, #P087.5, Carl Roth) at 4 °C. After fixation, the aortic tissue was stepwise dehydrated in ethanol followed by an incubation in Roti®Histol (Carl Roth) for 12 h. Afterwards the aortic tissue was neutralized and embedded in paraffin (#Carl Roth).

### 2.9 Murine platelet isolation and plasma preparation

Murine platelet isolation was conducted as described elsewhere (15). Briefly, blood was collected in 300 μL heparin solution (20 U/mL in PBS). Cell counts were determined using a hematology analyzer (Sysmex - KX21N, Norderstedt). The whole blood samples were centrifuged at 250x g for 5 min at RT. Platelet-rich plasma (PRP) was obtained by centrifugation at 50 x g for 6 min at RT. The PRP was washed twice in Tyrode’s buffer (136 mM NaCl, 0.4 mM Na2HPO4, 2.7 mM KCl, 12 mM NaHCO3, 0.1 % glucose, 0.35 % bovine serum albumin (BSA), pH 7.4), containing apyrase (0.02 U/mL) and prostacyclin (0.5 μM) by centrifugation at 650 x g for 5 min at RT. For experimental use, the isolated platelets were resuspended in Tyrode’s buffer supplemented with 1 mM CaCl_2_ (in absence of apyrase and prostacyclin). For plasma preparation whole blood was collected in heparin coated Microvettes® (Microvette® 500 Lithium heparin gel, #20.1346.100, Sarstedt) and centrifuged at 10,000 x g for 5 min at 4 °C. Plasma samples were stored at −70°C until use.

### 2.10 Preparation of murine platelet releasates

For the preparation of platelet releasates, murine platelets were isolated (40 × 10^6^) as described above and stimulated with CRP (0.1 or 1 μg/mL) or thrombin (0.1 U/mL) for 10 min at 37 °C. After stimulation sample were centrifuged at 650 x g for 5 min at 4 °C. The platelet releasates were collected and stored at −70°C before use.

### 2.11 Treatment of MHEC5-T cells with platelet releasates *in vitro*

The MHEC5-T (Mouse heart endothelial cell clone 5) cells were obtained from H. Langer, University Clinic Tübingen, Germany and cultured at 37°C in a humidified atmosphere with 5% CO_2_ in DMEM high glucose cell culture medium (#41965062, Gibco), containing 10% fetal calf serum (FCS; Life technologies), 1% penicillin-streptomycin (10,000 U/ml penicillin; 10,000 μg/ml streptomycin; Life technologies) and 200 mM L-glutamine (Life technologies). All following steps were conducted under sterile conditions. MHEC5-T cells were seeded in a density of 2.5 × 10^4^ cells/well into a 48-well plate and incubated with releasates of activated, respectively resting platelets for 3 h at 37 °C (5 % CO_2_). As controls, MHEC5-T were incubated with the indicated platelet agonists (negative control). Stimulation with 100 ng/mL TNFα served as positive control. After incubation, the cell culture supernatants were collected and cells were lysed using RLT Buffer from the RNeasy Mini Kit (#74106; Qiagen).

### 2.12 RNA isolation

RNA isolation of platelet releasate treated MHEC5-T cells was performed using the RNeasy Mini Kit (#74106; Qiagen). The isolation was conducted according to the manufacturer’s instructions. Afterwards RNA purity and concentration was analyzed using a spectro-photometer (BioPhotometer D30, Eppendorf).

### 2.13 cDNA synthesis

First, a DNA digestion was performed using DNAse I (#4716728001, Roche) to remove genomic DNA. Therefore, all samples were supplemented with DNAse I and incubated at 37 °C for 30 min. Reaction was stopped by incubating all samples at 75 °C for 10 min. Afterwards, cDNA synthesis was conducted by using the ImProm□IITM Reverse Transcription System (#A3800, Promega), following the manufacturer’s instructions. For cDNA synthesis, a total amount of 100 ng RNA was used.

### 2.14 Quantitative real-time PCR (qRT-PCR)

For analysing the endothelial gene expression of P-selectin and E-selectin after platelet releasate treatment, a quantitative real-time PCR was performed using the Fast Sybr Green Master Mix (#4385612, Thermo Fisher) by following the manufacturer’s protocol. The following primer were used: P-selectin (for: CATCTGGTTCAGTGCTTTGATCT; rev: ACCCGTGAGTTATTCCATGAGT) and E-selectin (for. ATGCCTCGCGCTTTCTCTC; rev: GTAGTCCCGCTGACAGTATGC). GAPDH (for: GGTGAAGGTCGGTGTGAACG; rev: CTCGCTCCTGGAAGATGGTG) served as housekeeping gene. The evaluation was performed according to the ΔΔCt method. Data were normalized to WT resting.

### 2.15 Immunofluorescence staining of murine aortic tissue

Paraffin embedded aortic tissue of the aneurysm segment 14 days after PPE was sliced in 5 μm sections using an automatic microtome (Microm HM355, Thermo Fisher Scientific). First, tissue sections were deparaffinised and hydrated. For antigen unmasking the tissue sections were heated up in citrate buffer (pH 6.0) for 10 min at 300 W. Afterwards, the tissue sections were blocked for 1 h at RT with protein blocking solution (#X0909, Dako). After washing with PBS the aortic tissue sections were specifically stained with primary antibodies for platelet GPIbα (CD42b #M042-0, Emfret Analytics, 1:50), respectively for neutrophils (Ly6G #551459; BD Pharming, 1:100). Incubation with primary antibodies was conducted overnight at 4 °C. Specific IgG primary antibodies served as controls. After washing with PBS the aortic tissue sections were incubated with a biotinylated secondary antibody (#BA-9400, Vector, 1:200) for 1h at RT. Hereafter, the tissue sections were incubated with a streptavidine FluorTM 660 conjugate (#50-4317-80, Thermo Fisher Scientific, 1:20) for 30 min at RT. Additionally, all sections were stained with DAPI (4′,6-Diamidine-2′-phenylindole dihydrochloride, #10236276001, Roche, 1:3000) to visualize cell nuclei within the aortic tissue. The stained tissue sections were embedded with mounting medium (#S3023, Dako) and stored at 4 °C until imaging. All images were generated using an Axio Observer. D1 microscope (Zeiss).

### 2.16 Flow cytometry

Flow cytometry was performed as described elsewhere (16). Briefly, heparinized murine whole blood was washed three times with Tyrode’s buffer by centrifugation at 650x g for 5 min. After washing the whole blood samples were diluted in Tyrode’s buffer containing 1 mM CaCl_2_. For the analysis of platelet activation, samples were stimulated with indicated agonists and specifically labeled with antibodies against P-selectin (CD62P, Wug.E9-FITC, #D200, Emfret Analytics) and active integrin αIIbβ3 (JON/A-PE, #D200; Emfret Analytics) in a ratio of 1:10 for 15 min at 37 °C. Reaction was stopped by adding 300 μL of PBS to all samples. To analyze glycoprotein (GP) expression, washed whole blood was labeled for GPIbα (CD42b, Xia.G5-PE, #M040-2 Emfret Analytics), GPVI (JAQ1-FITC, #M011-1, Emfret Analytics), integrin α5 (CD49e, Tap.A12-FITC, # M080-1, Emfret Analytics), or integrin β3 (CD61, Luc.H11-FITC, #M031-1, Emfret Analytics) in a ratio of 1:10 for 15 min at RT. For the analysis of platelet-neutrophil aggregate formation, washed whole blood was stimulated with the indicated agonists and labeled for the platelet marker GPIbα (CD42b, Xia.G5-PE, #M040-2 Emfret Analytics) and for the neutrophil marker Ly6G (1A8-APC, #127614, BD Biosciences) for 15 min at 37°C. For phosphatidylserine (PS)-exposure measurements, washed murine whole blood was diluted with binding buffer (containing 10 μM HEPES, 140 μM NaCl, 2.5 mM CaCl2, pH 7.4). Samples were stimulated with the indicated agonists and labelled with a PS detecting antibody (CyTM 5 Annexin V, #559934, BD Biosciences) for 15 min at RT. GPIbα (CD42b) was used as platelet specific marker. All samples were analyzed using a FACSCalibur flow cytometer (BD Biosciences).

### 2.17 Enzyme-Linked Immunosorbent Assay (ELISA)

All ELISA analysis were performed according to the manufacturer’s instructions. For quantification of Panx1 plasma concentration in AAA patients a human Panx1 ELISA (#39097, Signalway Antibody) was performed. For plasma analysis in mice, an IL-1β (#DY401, R&D Systems) and a MMP9 (#MMpT90, R&D Systems) ELISA were used.

### 2.18 Statistical analysis

Data are presented as arithmetic means ± SEM (Standard error of mean), statistical analysis was performed using GraphPad Prism 8 (version 8.4.3). Statistical differences were determined using a two-way ANOVA with a Sidak’s multiple comparison post-hoc test, an unpaired multiple t-test or an unpaired student’s t-test. Significant differences are indicated by asterisks (*** p < 0.001; ** p < 0.01; * p < 0.05).

## 3 Results

In this study, we analyzed plasma and platelets isolated from patients with AAA and platelet-specific Panx1 deficient mice in the PPE model that mimics human AAA pathology (17).

### 3.1 Enhanced Panx1 plasma levels but reduced phosphorylation of platelet Panx1 at Tyr^198^ in AAA patients

First, we analyzed the concentration of soluble Panx1 in the plasma of AAA patients. As shown in Figure 1A, increased amounts of soluble Panx1 was detected in patient’s plasma (Figure 1A). No alterations have been detected in the releasates of stimulated platelets using ADP and collagen-related peptide (CRP) to activate platelets and to induce the deletion of Panx1 from the platelet surface (Figure 1B). However, a significant increase in soluble Panx1 was only detected after platelet stimulation with 1 μg/mL CRP using platelets from AAA patients but not from healthy controls (Figure 1B). Next, we investigated the phosphorylation of Panx1 at Tyr^198^ as a marker for Panx1 activation. The activation of platelets from AAA patients resulted in elevated phosphorylation of Panx1 at Tyr^198^ only after stimulation with 1 μg/mL CRP but not with ADP or low dose CRP (0.1 μm/mL) (Figure 1C-D). However, reduced tyrosine phosphorylation was detected in platelets from AAA patients compared to healthy controls. However, total levels of Panx1 were comparable between both groups (Figure 1E).

**FIGURE 1.**
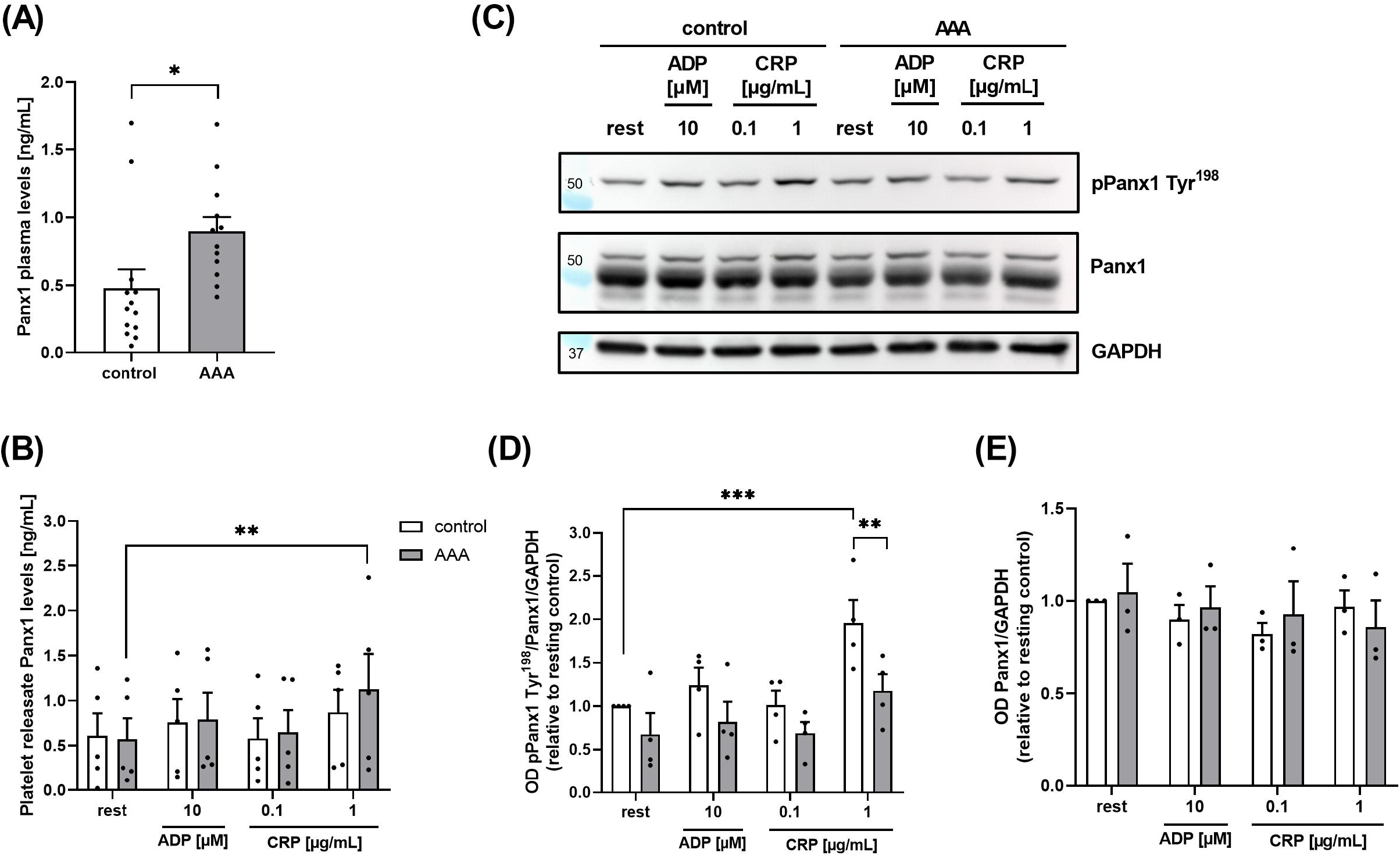
Soluble Panx1 plasma levels are enhanced in AAA patients, while platelet Panx1 Tyr^198^ phosphorylation is reduced. **(A)** Plasma concentration of soluble PANX1 in AAA patients (n=12), compared to age-matched controls (n=13). **(B)** Soluble Panx1 levels of platelet releasates from AAA patients (n=5) were analyzed after stimulation with indicated agonists (ADP [10 μM]; CRP [0.1/1 μg/mL]). Platelet releasates from age-matched controls served as controls (n=5). **(C)** Representative Western blot image and quantification (n=3-4). **(D-E)** Platelets of AAA patients and age-matched controls were isolated and stimulated with indicated agonists (ADP [10 μM]; CRP [0.1/1 μg/mL]). Platelet Panx1 protein content and tyrosine phosphorylation (pPanx1 Tyr^198^) were analyzed via Western blot. GAPDH served as loading control. Data are represented as mean values ± SEM. Statistical analysis was performed using **(A)** unpaired student’s t-test (**c-e**) two-way ANOVA with a Sidak’s multiple comparisons post-hoc test; *p < 0.05: **p < 0.01. AAA = Abdominal aortic aneurysm; ADP = Adenosine diphosphate; CRP = Collagen-related peptide; Panx1 = Pannexin-1.

### 3.2 Platelet-specific deletion of Panx1 leads to unaltered AAA formation and progression 28 days after PPE surgery

The results from AAA patients prompted us to analyze the impact of platelet Panx1 in further detail. We therefore induced experimental murine AAA utilizing the PPE model. We found that genetic loss of platelet-specific Panx1 did not alter diameter progression of AAA over a 28-day time course as shown by ultrasound tracking (Figure 2A-C). However, the incidence to develop AAA in mice was reduced in platelet-specific Panx1 knock-out mice (Figure 2D). Further, our data demonstrated reduced survival of *Panx1*^*fl/fl*^*-Pf4-Cre*^*+*^ mice after PPE surgery compared to controls (Figure 2E) potentially due to prolonged hemostasis in these mice (18). In line with unaltered aortic diameter expansion, we detected comparable aortic wall thickness between platelet-specific Panx1 knock-out and control mice as well as unaltered body weight (Figure 2F, Suppl.-Figure 1). Furthermore, MMP-9 levels in the plasma of platelet-specific Panx1 knock-out mice was only reduced by trend (Suppl.-Figure 1D). All these results suggest unaltered extracellular matrix (ECM) remodeling in these mice.

**FIGURE 2.**
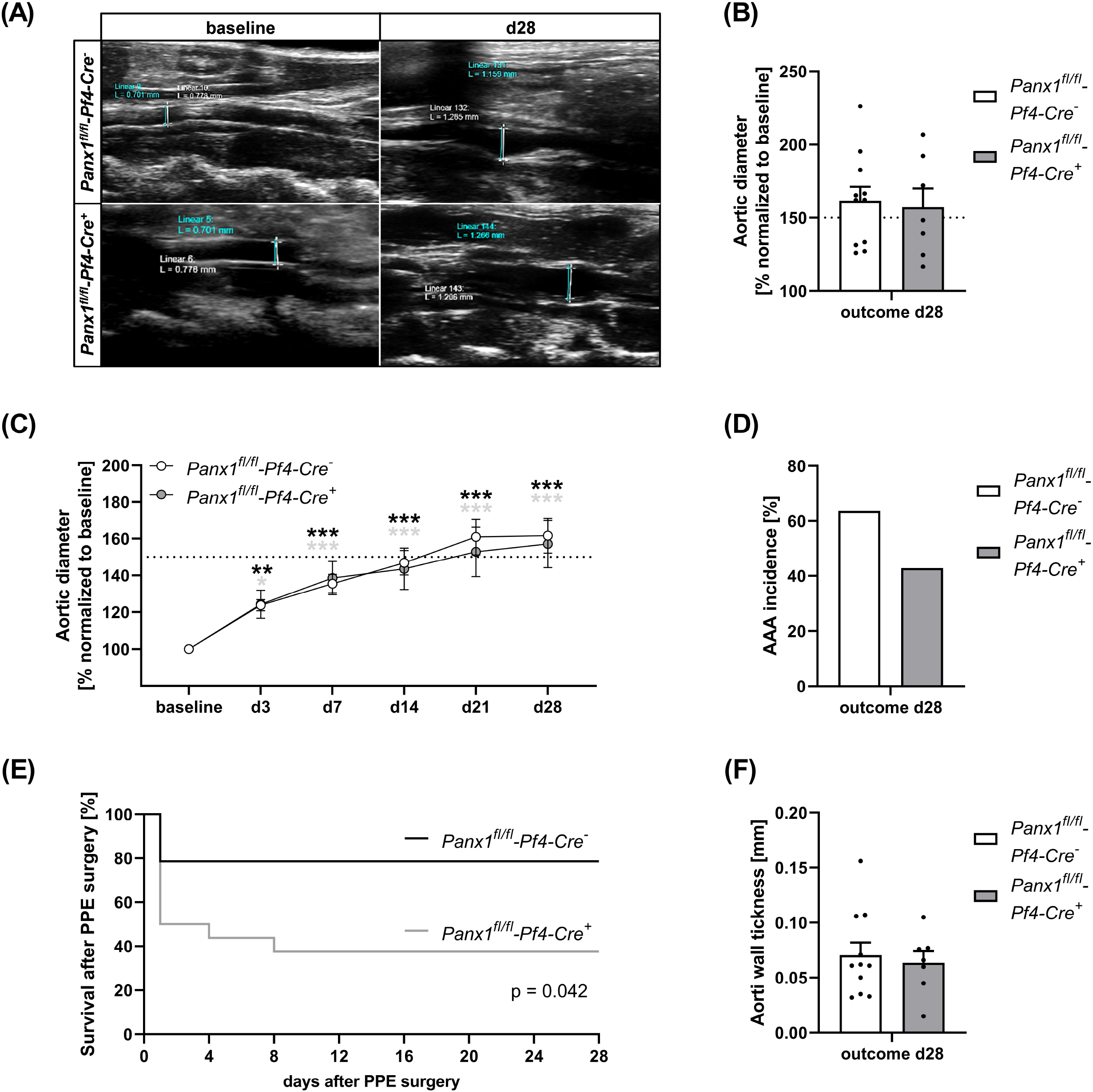
Platelet specific Panx1 deletion leads to unaltered AAA formation and progression 28 days after PPE surgery. **(A)** Representative Ultrasound images of the abdominal aorta from Panx1 WT (*Panx1*^*fl/fl*^*-Pf4-Cre*^*−*^; n=11) and Panx1 KO (*Panx1*^*fl/fl*^*-Pf4-Cre*^*+*^; n=7) mice at day 28 after PPE perfusion. **(B)** Aortic diameter in Panx1 WT (n=11) and Panx1 KO (n=7) mice at day 28 post PPE surgery. Data were normalized to baseline. **(C)** AAA development in Panx1 KO mice over a time period of 28 days after PPE (n=7), compared to Panx1 WT mice (n=11). Diameter progression of the abdominal aorta was measured via ultrasound imaging at day 3, 7, 14, 21 and 28, post PPE surgery. **(D)** AAA incidence in Panx1 WT and Panx1 KO mice at day 28 post PPE-surgery. Included are PPE operated mice that revealed an aortic diameter of ≥ 150 % 28 days after surgery. **(E)** Survival rate of Panx1 WT (n=11) and Panx1 KO (n=7) mice that underwent PPE surgery over a time period of 28 days. **(F)** Wall thickness within the aneurysm segment of Panx1 WT (n=11) and Panx1 KO (n=7) mice at day 28 after PPE. The aortic wall thickness was analyzed via ultrasound imaging. Data are represented as mean values ± SEM. Statistical analysis was performed using **(C)** two-way ANOVA with a Sidak’s multiple comparisons post-hoc test; indicated are statistical differences within one group **(D+F)** unpaired student’s t-test **(E)** Log-rank (Mantel-Cox) test; *p < 0.05, **p < 0.01, ***p < 0.001. AAA = Abdominal aortic aneurysm; Panx1 = Pannexin-1; PPE = Porcine pancreatic elastase perfusion.

### 3.3 Reduced platelet activation and pro-coagulant activity of platelets 14 and 28 days after PPE surgery

In a next step, we analyzed platelet activation after elastase infusion into the aortic wall using platelets from PPE mice at different time point after surgery (Figure 3). We did not detect major alterations in integrin αIIbβ3 (fibrinogen receptor) activation or P-selectin exposure as marker for degranulation as determined by flow cytometry (Suppl.-Fig. 2). In contrast, decreased Annexin-V binding as marker for pro-coagulant activity was detected at day 7 post PPE surgery following high doses of CRP that stimulates the major collagen receptor GPVI (Suppl.-Figure 2D). At 14 days after PPE surgery, we also detected decreased P-selectin exposure and active integrin αIIbβ3 in response to high doses of CRP and PAR4 peptide that stimulates the thrombin receptor PAR4 (Figure 3A). Defects in P-selectin exposure were even more pronounced after 28 days post-surgery, while defects in integrin activation have only been detected after stimulation of platelets with low dose of CRP (Figure 3B). Pro-coagulant activity was reduced with high dose of CRP but unaltered after thrombin stimulation of platelets (Figure 3B). However, GP exposure at the platelet surface was unaltered over the whole observation period of 28 days (Figure 3A-B, Suppl.-Figure 2C). In addition, we measured blood cell counts in naïve and PPE mice at different time points after surgery. Platelet and RBC counts were reduced on day 14 post-surgery while no alterations have been detected in mean platelet volume (MPV), WBC count and neutrophil-platelet conjugates after stimulation of platelets with either CRP or thrombin (Suppl.-Figure 3).

**FIGURE 3.**
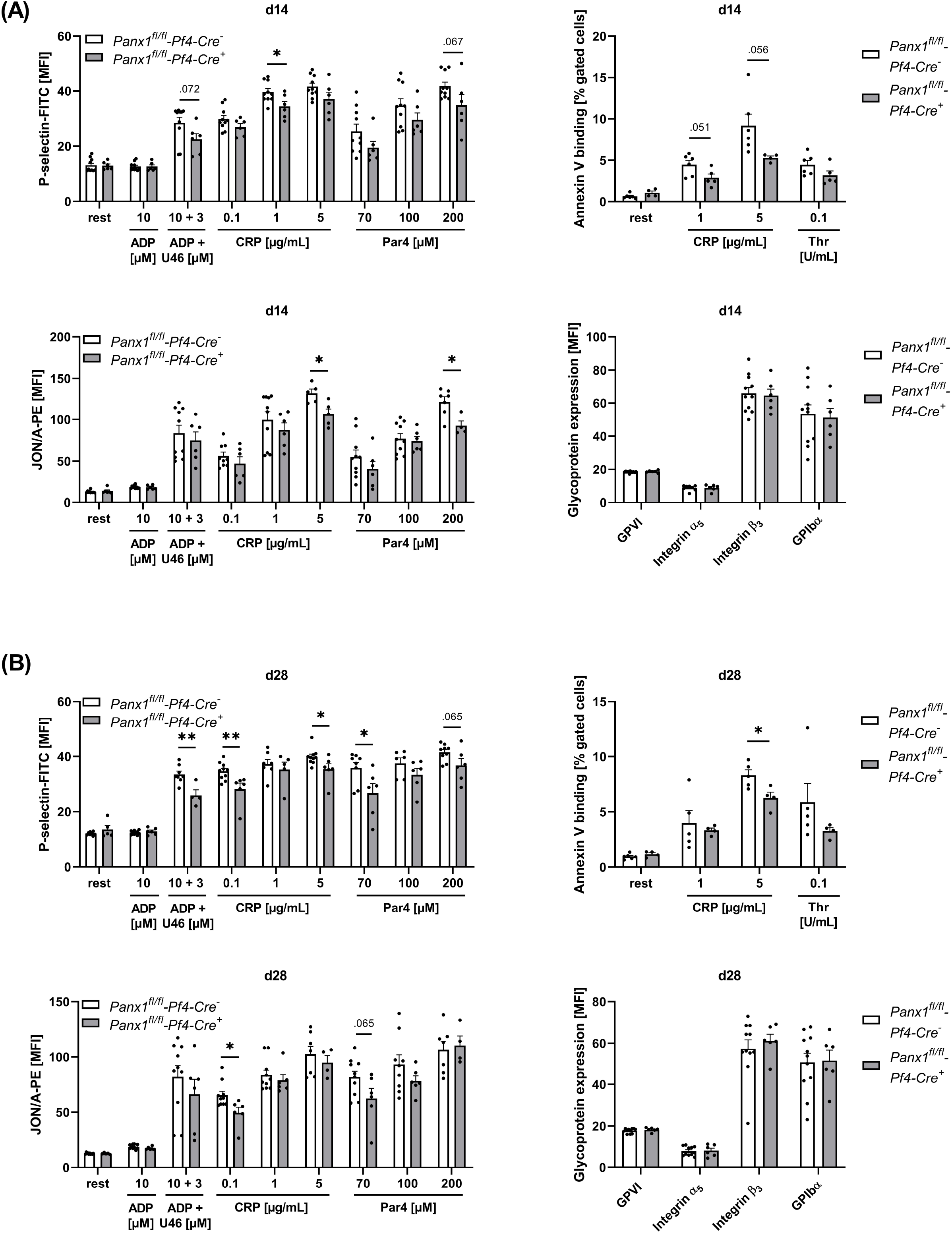
Platelets of Panx1 deficient mice reveal reduced platelet degranulation, activation and PS exposure during the remodeling phase of AAA development in the murine PPE model. Washed platelets from PPE operated Panx1 WT (*Panx1*^*fl/fl*^*-Pf4-Cre*^*−*^) and Panx1 KO (*Panx1*^*fl/fl*^*-Pf4-Cre*^*+*^) mice were analyzed via flow cytometry at day 14 **(A)**, and at day 28 **(B)** after surgery. **(A-B)** Platelet degranulation (P-selectin-FITC), integrin α_IIb_β_3_ activation (JON/A-PE) and PS-exposure (Annexin V-Cy5 binding) were analyzed after platelet stimulation with indicated agonists. In addition, platelet surface expression of GPVI, integrin α_5_, integrin β_3_ and GPIbα was analyzed via flow cytometry (PANX1 WT n=6-11; PANX1 KO n=4-6). Data are represented as mean values ± SEM. Statistical analysis was performed using an **(A-B)** unpaired multiple student’s t-test; *p < 0.05, **p < 0.01. ADP = Adenosine diphosphate; CRP = Collagen-related peptide; MFI = Mean fluorescence intensity; PANX1 = Pannexin-1; PAR4 = Protease-activated receptor 4 peptide; PPE = Porcine pancreatic elastase perfusion; Rest = resting; Thr = Thrombin; U46619 (U46) = Thromboxane A_2_ analogue.

### 3.4 Decreased platelet-mediated inflammation is characterized by reduced platelet-neutrophil conjugates and less cell migration into the aortic wall

To analyze the role of platelet Panx1 in inflammation, we analyzed Panx1 deficient platelets and their potential to form conjugates with neutrophils (Figure 4A-B). First, we analyzed the formation of platelet-neutrophil conjugates in the presence of LPS or TNF-α *in vitro* (Figure 4A). We detected significantly reduced platelet-neutrophil conjugates after stimulation using flow cytometry (Figure 4A). In a next step, we analyzed the formation of platelet-neutrophil conjugates in PPE mice at different time points. As shown in Figure 4B, reduced binding of neutrophils to platelets was detected after 14 days post-surgery. Reduced formation of platelet-neutrophil aggregates in platelet-specific Panx1 deficient mice resulted in reduced migration of neutrophils into the aortic wall of these mice 14 days after surgery (Figure 4C, Suppl.-Figure 5). Reduced migration of neutrophils was accompanied by reduced migration of platelets into aortic tissue 14 days after surgery that might be due to reduced platelet activation at this time point (Figure 3A, Figure 4D). However, the plasma level of the acute phase cytokine IL-1β was only reduced by trend in platelet-specific Panx1 knock-out mice (Suppl.-Figure 4A).

**FIGURE 4.**
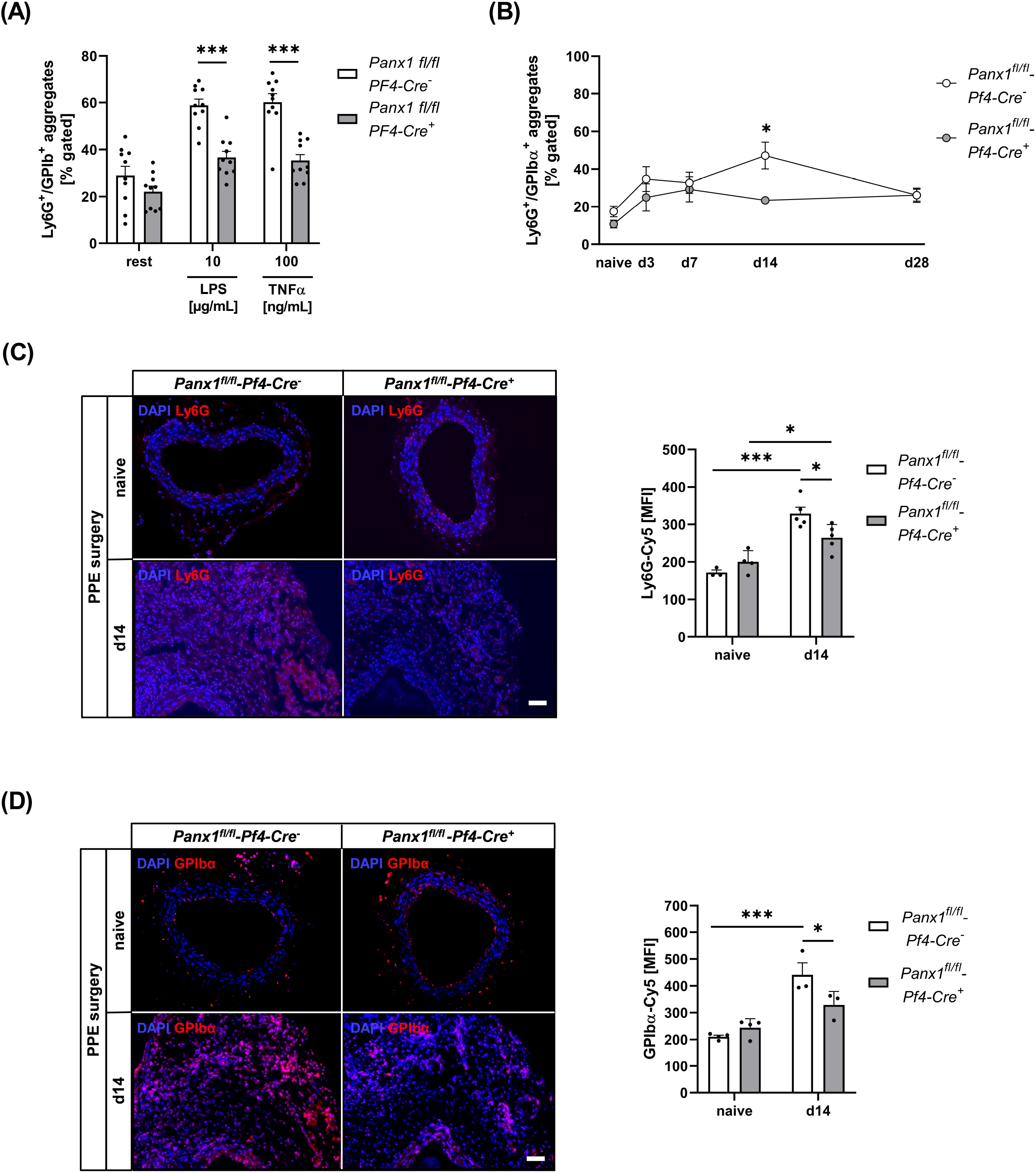
Reduced formation of platelet-neutrophil conjugates results in decreased migration of platelets and neutrophils into the vessel wall of Panx1 deficient mice 14 days after PPE surgery. **(A-B)** Platelet-neutrophil aggregates were analyzed via flow cytometry as double positive events for the platelet marker GPIbα (CD42b) and the neutrophil marker Ly6G. **(A)** Platelet-neutrophil aggregates in naïve Panx1 WT (*Panx1*^*fl/fl*^*-Pf4-Cre*^*−*^; n=10) and Panx1 KO (*Panx1*^*fl/fl*^*-Pf4-Cre*^*+*^; n=10) mice under resting conditions and after stimulation with LPS [10 μg/mL] or TNFα [100 ng/mL]. **(B)** Platelet-neutrophil aggregates in PPE operated Panx1 KO mice (n=6), compared to Panx1 WT mice (n=6-11). Aggregate formation was analyzed at day 0 (naive), 3, 7, 14 and 28 after PPE surgery. **(C)** Neutrophil migration into the aneurysm segment of PPE operated Panx1 WT (n= 5) and Panx1 KO (n=5) mice at day 14 post surgery (scale bar: 50 μm). Aortic tissue was specifically stained for neutrophils (Ly6G/Cy5; red) and DAPI (blue). Neutrophil migration was quantified as MFI of Ly6G positive cells within the aneurysm segment. Aortic tissue of naïve Panx1 WT (n=3) and Panx1 KO (n=4) mice served as control. **(D)** Migration of platelets into the aortic tissue of Panx1 WT (n=3) and Panx1 KO (n=3) mice at day 14 after PPE surgery (scale bar: 50 μm). Aortic tissue was specifically stained for platelets (GPIbα/Cy5; red) and DAPI (blue). Platelet migration into the vessel wall was quantified as MFI of GPIbα positive cells within the aneurysm segment. Aortic tissue of naïve Panx1 WT (n=3) and Panx1 KO (n=4) mice served as control. Data are represented as mean values ± SEM. Statistical analysis was performed using **(A)** unpaired multiple student’s t-test **(B, C-D)** two-way ANOVA with a Sidak’s multiple comparisons post-hoc test; *p < 0.05, ***p < 0.001. LPS = Lipopolysaccharide; MFI = Mean fluorescence intensity; Panx1 = Pannexin-1; PPE = Porcine Pancreatic Elastase perfusion; TNFα = Tumor necrosis factor alpha.

**FIGURE 5.**
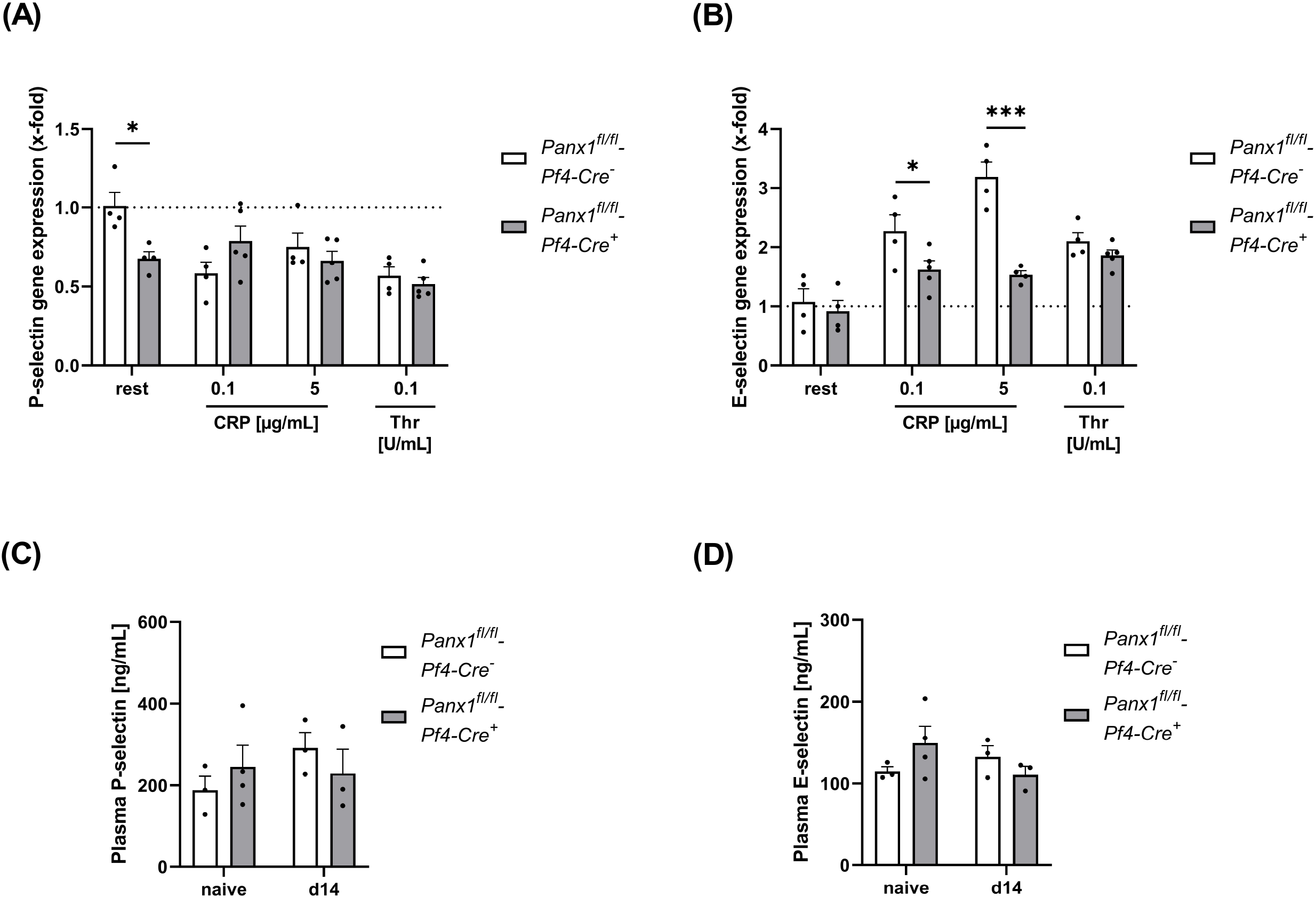
Platelet Panx1 modulates adhesion molecule gene expression in endothelial cells *in vitro*. **(A-B)** MHEC5-T cells (endothelial cell line) were incubated with platelet releasates of naïve Panx1 WT (*Panx1*^*fl/fl*^*-Pf4-Cre*^*−*^; n=4), respectively Panx1 KO (*Panx1*^*fl/fl*^*-Pf4-Cre*^*+*^; n=5) mice for 3h. For platelet releasates, isolated platelets were stimulated with the indicated agonists for 15 min beforehand. After incubation with platelet releasates for 3h, the MHEC5-T cells were harvested and gene expression of **(A)** P-selectin and **(B)** E-Selectin was analyzed via qRT-PCR. Data are normalised to WT rest. **(C-D)** Plasma concentration of soluble P-selectin and E-selectin of PPE operated Panx1 WT (*Panx1*^*fl/fl*^*-Pf4-Cre*^*−*^; n=3) and Panx1 KO (*Panx1*^*fl/fl*^*-Pf4-Cre*^*+*^; n=3) mice 14 days after surgery. Plasma samples from naïve Panx1 WT (n=3), respectively Panx1 KO (n=4) mice served as controls. Data are represented as mean values ± SEM. Statistical analysis was performed using **(A-D)** a two-way ANOVA with a Sidak’s multiple comparisons post-hoc test; *p < 0.05, ***p < 0.001. CRP = Collagen-related peptide; Panx1 = Pannexin-1; Rest = resting; Thr = Thrombin; PPE = Porcine Pancreatic Elastase perfusion.

### 3.5 Platelet Panx1 is important for the activation of endothelial cells

Platelet-mediated recruitment of inflammatory cells to inflamed tissue can be modulated by activated endothelial cells (19). To analyze, if platelet Panx1 modulates the activation of endothelial cells, we analyzed platelet-induced activation of MHEC5-T cells (endothelial cell line) *in vitro*. To this end, platelets were activated with CRP or thrombin and the supernatant was collected. Afterwards, we incubated MHEC5-T cells with platelet releasates of naïve Panx1 deficient and control mice for 3 h and analyzed gene expression of P-selectin and E-selectin in MHEC-5 T cells (Figure 5A-B). As shown in Figure 5A, the expression of P-selectin in MHEC5-T cells was only altered with resting but not with activated Panx1 deficient platelets (Figure 5A). In contrast, a strong reduction in E-selectin gene expression was detected when MHEC5-T cells were incubated with the supernatant of CRP-activated Panx1 deficient platelets compared to controls (Figure 5B). These results suggest that Panx1 in platelets is important for the activation of endothelial cells as reflected by gene expression of adhesion molecules such as P- and E-selectin (Figure 5A-B). To analyze plasma levels of soluble P-selectin and E-selectin in PPE mice, we used platelet-specific Panx1 deficient mice and the respective controls. As shown in Figure 5C-D, no alterations were detected 14 days after PPE infusion of mouse aorta between groups (Figure 5C-D).

## 4 Discussion

This study reveals enhanced Panx1 plasma levels as marker for enhanced Panx1 activation in AAA patients. In experimental AAA using the PPE model, a major contribution of platelet Panx1 channels in platelet activation, pro-coagulant activity of platelets and platelet-mediated inflammation has been detected. However, decreased platelet activation and inflammation in platelet-specific Panx1 knock-out mice did not affect ECM remodeling or wall thickness in PPE mice. Thus, AAA formation and progression was unaltered in theses mice as detected by aortic diameter expansion at different time points after intraluminal aortic elastase infusion.

Different studies in the past revealed a role of Panx1 channels for platelet activation and aggregation by releasing ATP using non-specific inhibitors named Probenecid (Prb) and Carbenoxolone (Cbx) and Panx1 knock-out mice (14, 20). A signaling mechanism was identified that includes collagen mediated GPVI activation leading to the phosphorylation of Src kinases resulting in ATP release via Panx1 channels into the extracellular space. This in turn amplifies P2×1 receptor activation and thus calcium flux important for platelet activation and aggregation. The activation of Panx1 channels by Src kinases is crucial for the phosphorylation of Panx1 at Tyr^198^ and Tyr^308^, suggesting that Src kinases are key regulators in Panx1 activation in platelets (13, 18). Beside GPVI mediated activation of Panx1, G protein-coupled receptor activation and thromboxane mediated signaling pathways are able to amplify Panx1 phosphorylation (13). Panx1 activation and subsequent non-vesicular ATP release from platelets amplifies thrombus formation and therefore plays an important role in hemostasis and thrombosis (13, 18).

Due to the important role of extracellular ATP as inflammatory molecule (21, 22), a role for Panx1 in inflammation was discovered by different groups. It was shown that Panx1 dependent NLRP3 inflammasome activation results in IL-1β and IL-18 production, leading to toll-like receptor (TLR) signaling and the stimulation of purinergic receptors by high extracellular ATP concentrations (22). The study of Lohman and colleagues revealed that controlled ATP release via Panx1 channels supports intercellular communication and regulates leukocyte emigration into venous endothelium during acute inflammation (23). Furthermore, Panx1 channels were identified as mediators of neuroinflammation (11) and act as drives of inflammation in acute myocardial infarction (24). An important role for Panx1 in inflammation has been observed especially for endothelial cells, because Panx1 at the plasma membrane of these cells mediates vascular inflammation during lung ischemia-reperfusion injury (25) and regulates cardiac responses to acute myocardial infarction (26). Thus, it is not surprising that endothelial cell specific Panx1 deficiency but not smooth muscle cell specific Panx1 deficiency resulted in reduced AAA formation compared to respective WT mice in the topical PPE model (12). The authors detected reduced acute phase cytokines such as HMGB1, IL-17, MCP1 and TNFα in Panx1 deficient mice compared to elastase-treated control mice. Additionally, a reduced number of infiltrating macrophages, neutrophils and CD3+ T cells were found in elastase-treated endothelial cell specific Panx1 deficient mice. These mice displayed decreased elastic fibre disruption compared to their respective controls. Application of the Panx1 inhibitor Prb to PPE mice inhibited leukocyte transmigration, aortic inflammation and remodeling to attenuate AAA formation (12). All these data suggest a major contribution of Panx1 channels in inflammation and AAA formation with a dominant role of endothelial Panx1 in these processes. However, we did not observe differences in aortic diameters following platelet-specific Panx1 knock-out mice using the PPE infusion model. This may be explained by destruction of the endothelial cell layer due to enzymatic degradation and constant pressure during the procedure, while the topical elastase model is characterized by degradation of the vessel wall starting at the adventitial layer, leaving the endothelium initially unaffected. Therefore, platelet-specific Panx1 mediated activation of endothelial cells cannot be observed in the PPE model and might explain unaltered P- and E-selectin plasma levels in PPE mice of both groups.

The here presented results amplify the important role of Panx1 for the inflammatory response in CVDs/atherosclerotic diseases and especially in AAA; and provide evidence for the first time that platelet Panx1 is involved in these processes. Since many years, a role for platelets in acute and chronic inflammation in CVDs, sepsis, cancer, and in severe neurological disorders such as multiple sclerosis or stroke has been established (19, 27–29). In experimental AAA, Clopidogrel treatment of Ang-II infused mice reduced vascular inflammation and AAA progression (30). Treatment of xenografted rats with Abciximab limited aneurysm expansion (31). Using the PPE mouse model, Hannawa and colleagues identified P-selectin as important modulator of inflammation and aortic wall remodeling (32). However, the authors used a P-selectin knock-out mouse model that disrupts P-selectin exposure in endothelial cells and in platelets. Thus, further experiments are needed to analyze the independent contribution of endothelial cells and platelets in experimental AAA formation and progression. Our results with platelet specific Panx1 knock-out mice revealed that also Panx1 channels at the platelet surface trigger inflammation in AAA. However, aortic wall remodeling was not altered in these mice suggesting that platelet Panx1 is implicated solely in the inflammatory response but not in ECM remodeling in AAA in contrast to endothelial Panx1 that modulates both processes. Furthermore, reduced inflammation did not result in reduced AAA formation as shown by unaltered aortic diameter expansion and wall thickness in platelet-specific Panx1 knock-out mice compared to controls.

In summary, this study adds important knowledge about the role of platelets and Panx1 channels on the formation and progression of AAA. Identification of the defined role of platelets in AAA may help to identify novel therapeutic targets for the prevention of AAA formation and progression because the efficacy and safety of antiplatelet therapy has not been established in patients with AAA to date.

## Supporting information

Metz Feige et al_Supplementary Material

## 5 Data Availability Statement

The original contributions presented in the study are included in the article, further inquiries can be directed to the corresponding author.

## 7 Author Contributions

HS and ME designed the study. LMM, TF, LB, AE, JM, LMT, and FR performed experiments. HS, CQ, NG, MK and ME analyzed and interpreted data. TF and ME wrote the manuscript with all authors providing feedback. MK provided infrastructure. LMM and TF contributed equally to this study.

## 9 Acknowledgments

We thank Martina Spelleken for excellent technical assistance and Julia Odendahl for PPE surgery.

## References

1. Aziz F, Kuivaniemi H. Role of matrix metalloproteinase inhibitors in preventing abdominal aortic aneurysm. Annals of vascular surgery. 2007;21(3):392–401.

2. Busch A, Chernogubova E, Jin H, Meurer F, Eckstein HH, Kim M, et al. Four Surgical Modifications to the Classic Elastase Perfusion Aneurysm Model Enable Haemodynamic Alterations and Extended Elastase Perfusion. European journal of vascular and endovascular surgery : the official journal of the European Society for Vascular Surgery. 2018;56(1):102–9.

3. Ullery BW, Hallett RL, Fleischmann D. Epidemiology and contemporary management of abdominal aortic aneurysms. Abdominal radiology (New York). 2018;43(5):1032–43.

4. Vardulaki KA, Walker NM, Day NE, Duffy SW, Ashton HA, Scott RA. Quantifying the risks of hypertension, age, sex and smoking in patients with abdominal aortic aneurysm. The British journal of surgery. 2000;87(2):195–200.

5. Lindquist Liljeqvist M, Hultgren R, Bergman O, Villard C, Kronqvist M, Eriksson P, et al. Tunica-Specific Transcriptome of Abdominal Aortic Aneurysm and the Effect of Intraluminal Thrombus, Smoking, and Diameter Growth Rate. Arteriosclerosis, thrombosis, and vascular biology. 2020;40(11):2700–13.

6. Sakalihasan N, Michel JB, Katsargyris A, Kuivaniemi H, Defraigne JO, Nchimi A, et al. Abdominal aortic aneurysms. Nature reviews Disease primers. 2018;4(1):34.

7. Shimizu K, Mitchell RN, Libby P. Inflammation and cellular immune responses in abdominal aortic aneurysms. Arteriosclerosis, thrombosis, and vascular biology. 2006;26(5):987–94.

8. Bao L, Locovei S, Dahl G. Pannexin membrane channels are mechanosensitive conduits for ATP. FEBS letters. 2004;572(1-3):65–8.

9. Sandilos JK, Chiu YH, Chekeni FB, Armstrong AJ, Walk SF, Ravichandran KS, et al. Pannexin 1, an ATP release channel, is activated by caspase cleavage of its pore-associated C-terminal autoinhibitory region. The Journal of biological chemistry. 2012;287(14):11303–11.

10. Abitbol JM, O’Donnell BL, Wakefield CB, Jewlal E, Kelly JJ, Barr K, et al. Double deletion of Panx1 and Panx3 affects skin and bone but not hearing. Journal of molecular medicine (Berlin, Germany). 2019;97(5):723–36.

11. Seo JH, Dalal MS, Contreras JE. Pannexin-1 Channels as Mediators of Neuroinflammation. International journal of molecular sciences. 2021;22(10).

12. Filiberto AC, Spinosa MD, Elder CT, Su G, Leroy V, Ladd Z, et al. Endothelial pannexin-1 channels modulate macrophage and smooth muscle cell activation in abdominal aortic aneurysm formation. Nature communications. 2022;13(1):1521.

13. Metz LM, Elvers M. Pannexin-1 Activation by Phosphorylation Is Crucial for Platelet Aggregation and Thrombus Formation. International journal of molecular sciences. 2022;23(9).

14. Taylor KA, Wright JR, Vial C, Evans RJ, Mahaut-Smith MP. Amplification of human platelet activation by surface pannexin-1 channels. Journal of thrombosis and haemostasis : JTH. 2014;12(6):987–98.

15. Donner L, Toska LM, Krüger I, Gröniger S, Barroso R, Burleigh A, et al. The collagen receptor glycoprotein VI promotes platelet-mediated aggregation of β-amyloid. Science signaling. 2020;13(643).

16. Klatt C, Krüger I, Zey S, Krott KJ, Spelleken M, Gowert NS, et al. Platelet-RBC interaction mediated by FasL/FasR induces procoagulant activity important for thrombosis. The Journal of clinical investigation. 2018;128(9):3906–25.

17. Busch A, Bleichert S, Ibrahim N, Wortmann M, Eckstein HH, Brostjan C, et al. Translating mouse models of abdominal aortic aneurysm to the translational needs of vascular surgery. JVS-vascular science. 2021;2:219–34.

18. Molica F, Meens MJ, Pelli G, Hautefort A, Emre Y, Imhof BA, et al. Selective inhibition of Panx1 channels decreases hemostasis and thrombosis in vivo. Thrombosis research. 2019;183:56–62.

19. Ed Rainger G, Chimen M, Harrison MJ, Yates CM, Harrison P, Watson SP, et al. The role of platelets in the recruitment of leukocytes during vascular disease. Platelets. 2015;26(6):507–20.

20. Molica F, Morel S, Meens MJ, Denis JF, Bradfield PF, Penuela S, et al. Functional role of a polymorphism in the Pannexin1 gene in collagen-induced platelet aggregation. Thrombosis and haemostasis. 2015;114(2):325–36.

21. Corriden R, Insel PA. New insights regarding the regulation of chemotaxis by nucleotides, adenosine, and their receptors. Purinergic signalling. 2012;8(3):587–98.

22. Gombault A, Baron L, Couillin I. ATP release and purinergic signaling in NLRP3 inflammasome activation. Frontiers in immunology. 2012;3:414.

23. Lohman AW, Leskov IL, Butcher JT, Johnstone SR, Stokes TA, Begandt D, et al. Pannexin 1 channels regulate leukocyte emigration through the venous endothelium during acute inflammation. Nature communications. 2015;6:7965.

24. Koval M, Cwiek A, Carr T, Good ME, Lohman AW, Isakson BE. Pannexin 1 as a driver of inflammation and ischemia-reperfusion injury. Purinergic signalling. 2021;17(4):521–31.

25. Sharma AK, Charles EJ, Zhao Y, Narahari AK, Baderdinni PK, Good ME, et al. Pannexin-1 channels on endothelial cells mediate vascular inflammation during lung ischemia-reperfusion injury. American journal of physiology Lung cellular and molecular physiology. 2018;315(2):L301–l12.

26. Good ME, Young AP, Wolpe AG, Ma M, Hall PJ, Duffy CK, et al. Endothelial Pannexin 1 Regulates Cardiac Response to Myocardial Infarction. Circulation research. 2021;128(8):1211–3.

27. Koupenova M, Clancy L, Corkrey HA, Freedman JE. Circulating Platelets as Mediators of Immunity, Inflammation, and Thrombosis. Circ Res. 2018;122(2):337–51.

28. Rawish E, Nording H, Münte T, Langer HF. Platelets as Mediators of Neuroinflammation and Thrombosis. Front Immunol. 2020;11:548631.

29. Thomas MR, Storey RF. The role of platelets in inflammation. Thrombosis and haemostasis. 2015;114(3):449–58.

30. Liu O, Jia L, Liu X, Wang Y, Wang X, Qin Y, et al. Clopidogrel, a platelet P2Y12 receptor inhibitor, reduces vascular inflammation and angiotensin II induced-abdominal aortic aneurysm progression. PloS one. 2012;7(12):e51707.

31. Touat Z, Ollivier V, Dai J, Huisse MG, Bezeaud A, Sebbag U, et al. Renewal of mural thrombus releases plasma markers and is involved in aortic abdominal aneurysm evolution. The American journal of pathology. 2006;168(3):1022–30.

32. Hannawa KK, Cho BS, Sinha I, Roelofs KJ, Myers DD, Wakefield TJ, et al. Attenuation of experimental aortic aneurysm formation in P-selectin knockout mice. Annals of the New York Academy of Sciences. 2006;1085:353–9.

