## Supplementary material for "Platelet pannexin-1 channels modulate inflammation during abdominal aortic aneurysm formation": Metz Feige et al_Supplementary Material

### 1 Supplementary Figures and Tables

#### 1.1 Supplementary Figures

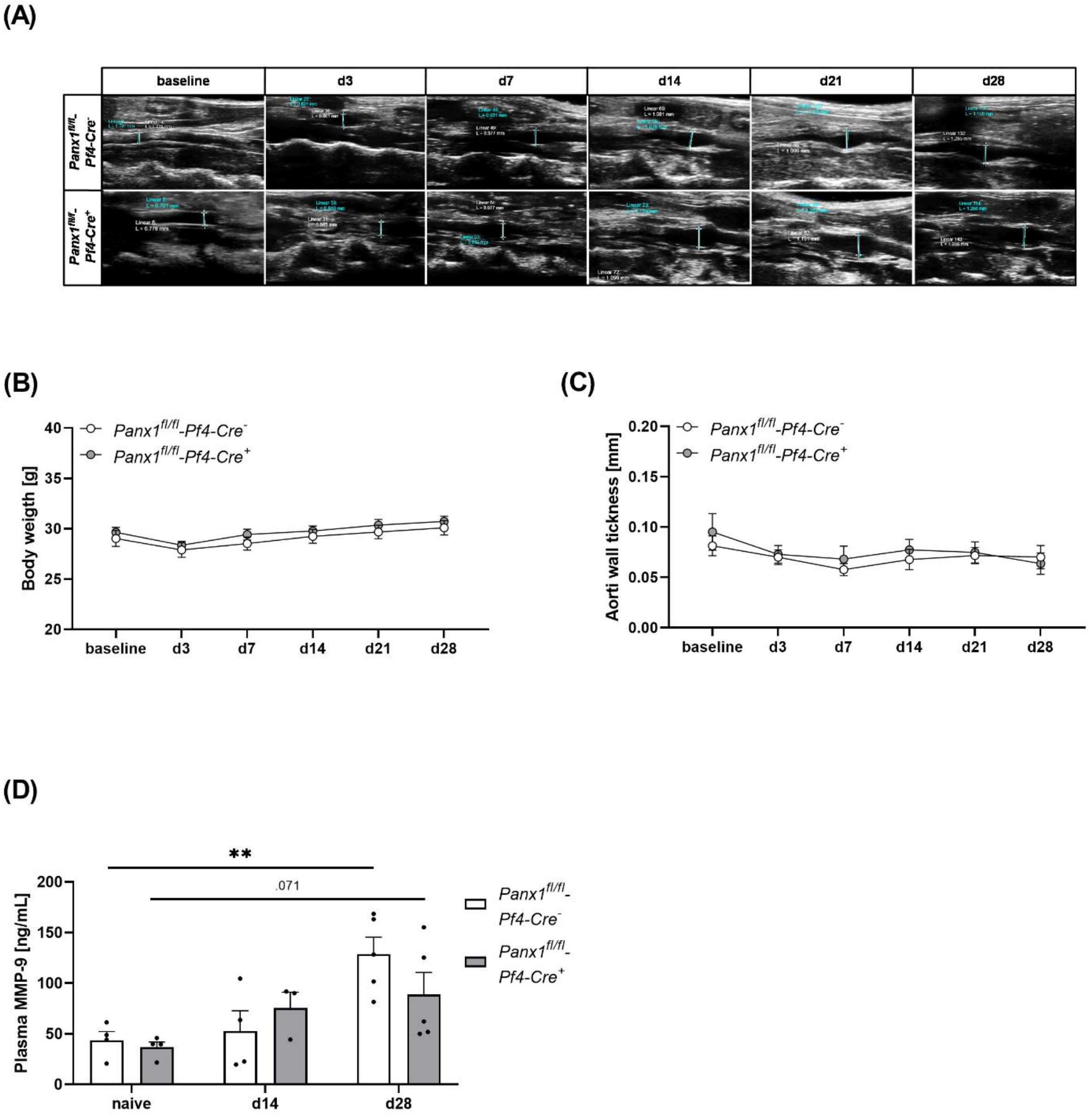


**Supplementary Figure 1.** Platelet specific Panx1 deletion leads to unaltered aortic wall thickness during AAA development after PPE surgery. **(A)** Representative Ultrasound images of the abdominal aorta from Panx1 WT (Panx1 fl/fl PF4-Cre$^{-}$; n=11) and Panx1 KO (Panx1 fl/fl PF4-Cre$^{+}$; n=7) mice at day 28 after PPE perfusion. Ultrasound imaging was conducted day 0 (pre surgery) and at day 3, 7, 14,

21 and 28 after PPE. **(B)** Body weight of Panx1 WT (n=11) and Panx1 KO (n=7) mice during PPE induced AAA formation. **(C)** Aortic wall thickness progression within the aneurysm segment in Panx1 WT (n=11) and Panx1 KO (n=7) mice over a time period of 28 days after PPE. The aortic wall thickness was determined via ultrasound measurements at the indicated time points; **(D)** MMP9 plasma concentration of PPE operated Panx1 KO mice (n=3-5) at day 14 and day 28, compared to Panx1 WT mice (n=3-5). Plasma of naïve Panx1 WT (n=4) and Panx1 KO (n=4) mice served as control. Data are represented as mean values ± SEM. Statistical analysis was performed using two-way ANOVA with a Sidak's multiple comparisons post-hoc test. AAA = Abdominal aortic aneurysm; MMP-9 = Matrix metalloproteinase-9; Panx1 = Pannexin-1; PPE = Porcine pancreatic elastase perfusion.

.

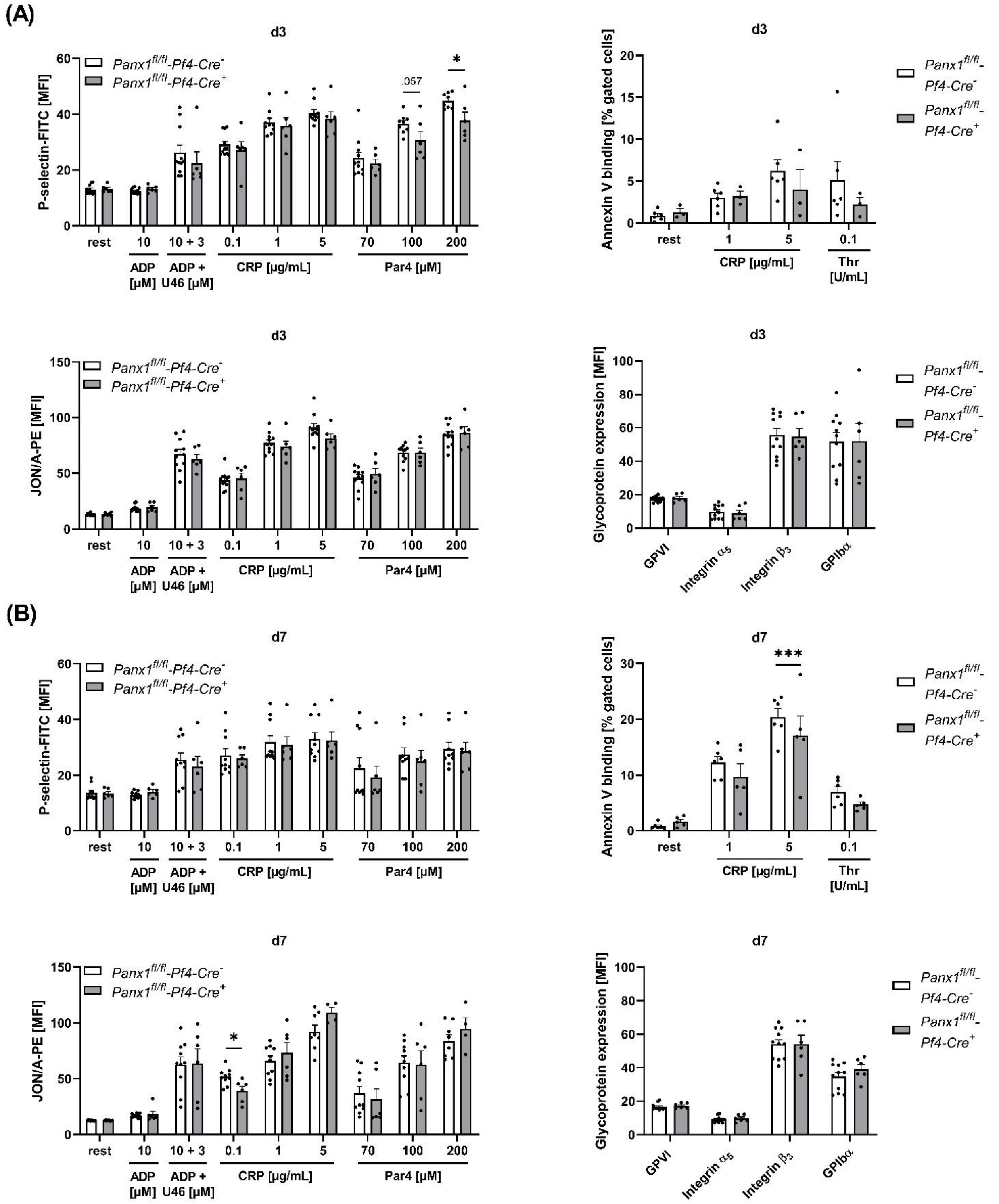


**Supplementary Figure 2.** Platelets of Panx1 deficient mice reveal an unaltered platelet degranulation, activation and PS-exposure during the inflammatory phase of AAA development. Washed platelets from PPE operated Panx1 WT (*Panx1 fl/fl PF4-Cre*$^{-}$) and Panx1 KO (*Panx1 fl/fl PF4-Cre*$^{+}$) mice were analysed via flow cytometry at day 3 **(A)**, and at day 7 **(B)** after surgery. **(A-B)** Platelet degranulation (P-selectin-FITC), integrin $\alpha_{IIb}\beta_3$ activation (JON/A-PE) and PS-exposure (Annexin V-Cy5 binding) were analysed after platelet stimulation with indicate agonists. In addition, platelet surface expression of GPVI, integrin $\alpha_5$, integrin $\beta_3$ and GPIbα was analysed via flow cytometry (Panx1 WT n=5-11; Panx1 KO n=4-6). Data are represented as mean values ± SEM. Statistical analysis was performed using an **(A-B)** unpaired multiple student's t-test; $*p < 0.05$, $***p < 0.001$. ADP = Adenosine diphosphate; CRP = Collagen-related peptide; MFI = Mean fluorescence intensity; Panx1 = Pannexin-1; PAR4 = Protease-activated receptor 4 peptide; PPE = Porcine pancreatic elastase perfusion; Rest = resting; Thr = Thrombin; U46619 (U46) = Thromboxane $A_2$ analogue.

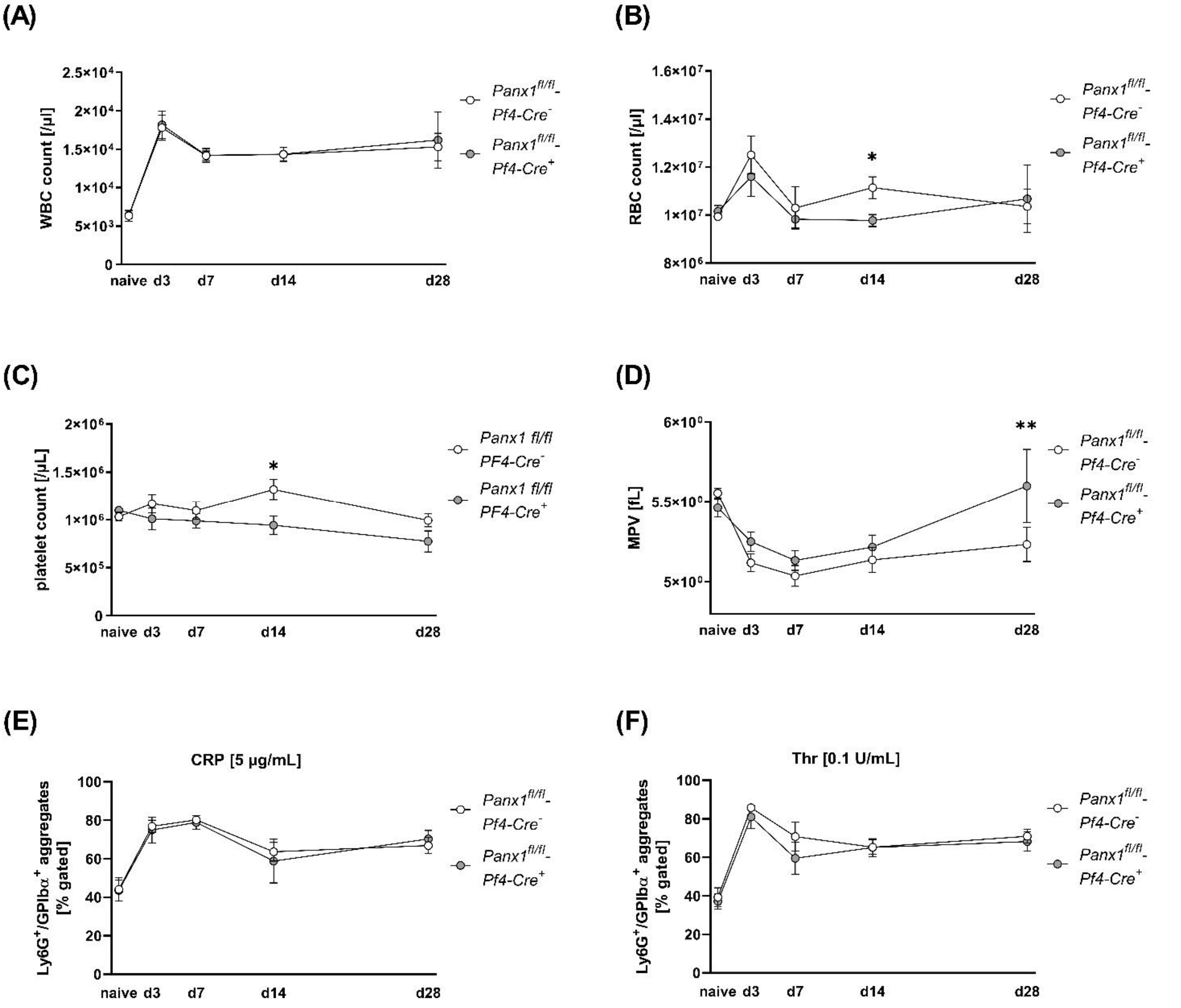


**Supplementary Figure 3.** Platelet Panx1 deficient mice reveal alteration in blood cell count and MVP during PPE induced AAA formation**. (A)** WBC, **(B)** RBC, **(C)** platelet counts and **(D)** the MPV of Panx1 WT (*Panx1 fl/fl PF4-Cre*$^{-}$; n=7-11), respectively Panx1 KO (*Panx1 fl/fl PF4-Cre*$^{+}$; n=5-7) mice were analysed at day 0 (naïve), 3, 7, 14, and day 28 after PPE surgery. **(E-F)** Platelet neutrophil aggregates in PPE operated Panx1 WT (n=6-11) and Panx1 KO (n=4-6) mice after stimulation with **(E)** CRP [5 µg/mL] or **(F)** thrombin [0.1 U/mL]. Aggregate formation was analysed at day 0 (naive), 3, 7, 14 and 28 after PPE surgery via flow cytometry as double positive events for the platelet marker GPIbα (CD42b) and the neutrophil marker Ly6G. Data are represented as mean values ± SEM. Statistical analysis **(A-F)** was performed using a two-way ANOVA with a Sidak's multiple comparisons post-hoc test; *$p < 0.05$, **$p < 0.01$. CRP = Collagen-related peptide; MPV = Mean platelet volume; Panx1 = Pannexin-1; PPE = Porcine pancreatic elastase perfusion; RBCs = Red Blood Cells, Thr = Thrombin; WBCs = White Blood Cells.

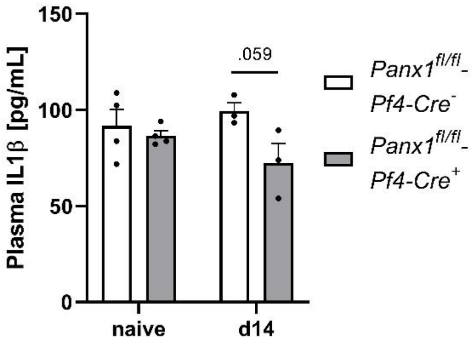


**Supplementary Figure 4.** Plasma IL1β level are unaltered in platelet Panx1 deficient mice during PPE induced AAA progression. IL1β plasma concentration of PPE operated Panx1 KO mice (n=3-5) at day 14 post surgery, compared to Panx1 WT mice (n=3-5). Plasma of naïve Panx1 WT (n=4) and Panx1 KO (n=4) mice served as control. Data are represented as mean values ± SEM. Statistical analysis was performed using a two-way ANOVA with a Sidak's multiple comparisons post-hoc test; *$p < 0.05$, **$p < 0.01$, ***$p < 0.001$. IL-1β = Interleukin-1β; Panx1 = Pannexin-1; Panx1 = Pannexin-1; PPE = Porcine Pancreatic Elastase.

**(A)**

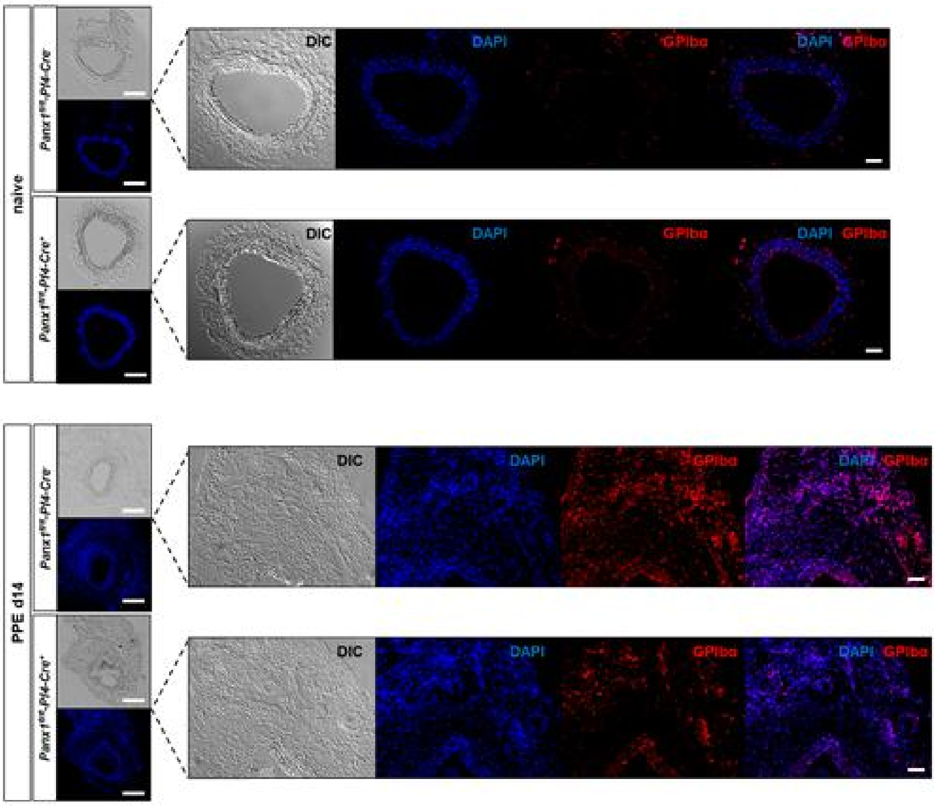
naive
PPE d14
DIC
DAPI
GPIbα
DAPI GPIbα

**(B)**

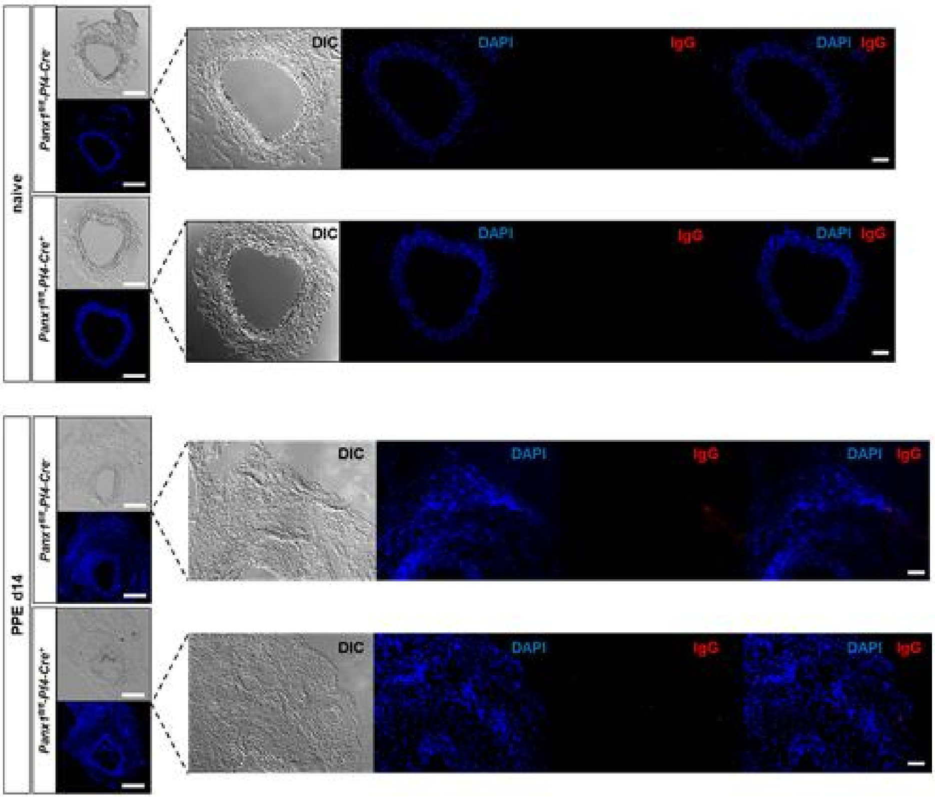
naive
PPE d14
DIC
DAPI
IgG
DAPI IgG

**Supplementary Figure 5.** Immunofluorescence staining of platelets within the aortic tissue of PPE operated Panx1 deficient mice. **(A)** Immunofluorescence staining of platelets (GPIbα/Cy5; red) in the aortic tissue of Panx1 WT (*Panx1 fl/fl PF4-Cre*$^{-}$; n=3) and Panx1 KO (*Panx1 fl/fl PF4-Cre*$^{+}$; n=3) mice at day 14 after PPE surgery and **(B)** the respective IgG controls (scale bar: 200 µm and 50 µm). Aortic tissue of naïve Panx1 WT (n=3) and Panx1 KO (n=4) mice served as control. DIC= Differential interference contrast; Panx1 = Pannexin-1

(A)

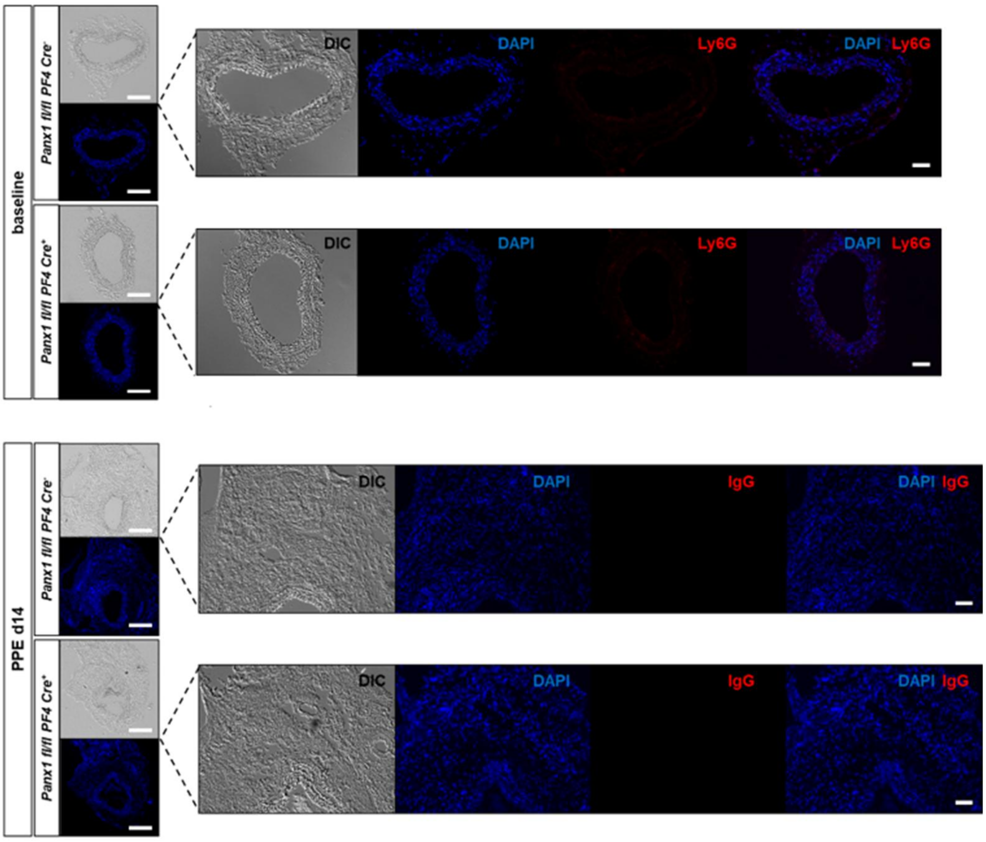
baseline
Panx1 fl/fl PF4 Cre-
Panx1 fl/fl PF4 Cre+
DIC
DAPI
Ly6G
DAPI Ly6G
PPE d14
IgG
DAPI IgG

(B)

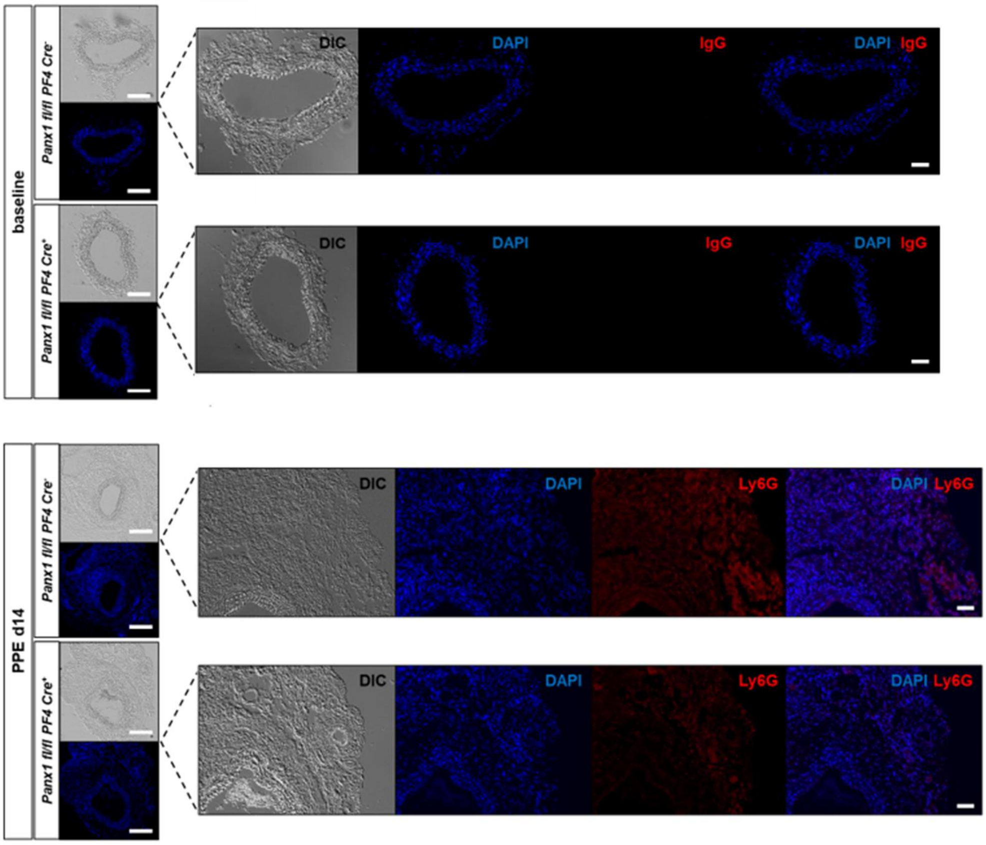
baseline
Panx1 fl/fl PF4 Cre-
Panx1 fl/fl PF4 Cre+
DIC
DAPI
IgG
DAPI IgG
PPE d14
Ly6G
DAPI Ly6G

**Supplementary Figure 6.** Immunofluorescence staining of neutrophils within the aortic tissue of PPE operated Panx1 deficient mice. **(A)** Immunofluorescence staining of neutrophils (Ly6G/Cy5; red) in the aortic tissue of Panx1 WT (*Panx1 fl/fl PF4-Cre*$^{-}$; n=3) and Panx1 KO (*Panx1 fl/fl PF4-Cre*$^{+}$; n=3) mice at day 14 after PPE surgery and **(B)** the respective IgG controls (scale bar: 200 µm and 50 µm). Aortic tissue of naïve Panx1 WT (n=3) and Panx1 KO (n=4) mice served as control. DIC= Differential interference contrast; Panx1 = Pannexin-1
